# Multilamellar mesoporous silica nanoparticles using a cationic co-surfactant dual-templating method

**DOI:** 10.1101/2021.02.25.432869

**Authors:** Timo Froyen, Niels Geysmans, Ulrique Vounckx, An Hardy

**Affiliations:** DESINe, Hasselt University, Institute for Materials Research (IMO), and imec division imomec, Agoralaan, 3590 Diepenbeek, Belgium research group, Biomedical Research Institute

**Keywords:** multilamellar, mesoporous, vesicular, worm-like, dual-templating

## Abstract

The utility of mesoporous silica nanoparticles (MSNs) has been repeatedly proven in a wide range of biomedical applications. The general morphology of these particles is easily modifiable by various post-grafting possibilities and adjustments within the surfactant-based template. The synthesis of multilamellar vesicular silica nanoparticles has led to the discovery of beneficial attributes regarding said particles. Depending on the synthesis process, various parameters are affected including packaging capacity, stability, drug adsorption and release. This research focused on synthesis and characterization of multilamellar MSNs using a cationic-cationic co-surfactant templating route testing various ratios of cetyltrimethylammonium bromide (CTAB) and didodecyldimethylammonium bromide (DDAB). TEM imaging showed clear differences in size and morphology between the different samples, and was further characterized by BET and BJH analysis. All multilamellar nanoparticles did exhibit a similar pore size distribution and overall gradual release of drug contents. However, the degree of drug adsorption and overtime drug release was clearly influenced by the number of layers of the MSNs, proving the utility of adjusting the template. Further experiments could be conducted to validate the utility of beta-cyclodextrin as a template regulator and to investigate both biocompatibility and biodegradability of the multilamellar MSNs.

## 1. INTRODUCTION

Gradual hydrolysis and water- or alcohol condensation of silica precursors (SiO2) are responsible for the spontaneous self-assembly of mesoporous silica nanospheres (MSNs) under the right conditions and is often referred to as a Stöber-like synthesis (1). These easily fabricated structures possess various properties, making them a popular concept for a multitude of nanotechnological fields. Biomedical applications involving diagnostic imaging, drug/gene delivery and tissue regeneration are highly advancing in MSN optimization, but also other applications such as confined-space catalysis, antireflective coatings, adsorption, polymer filling and optical devices have found utility in nano-scaled mesoporous silica structures (2–7). The popularity revolving MSNs can be attributed to numerous advantageous characteristics: (1) a large specific surface area generated by porosity, opposed to non-porous silica material, (2) a controllable pore size depending on the amount of silica precursors and surfactant, (3) a controllable pore volume depending on alkyl chain length and swelling agents, (4) a narrow pore size distribution, contributing to the uniformity, (5) good biocompatibility, which combined with (6) good biodegradability, avoids any adverse effects potentially caused by the immune system. By tuning the pore size within the meso-range (2 nm to 50 nm), uptake of proteins, enzymes or drug molecules is favored and consequently affects eventual release of said molecules (8). An interesting alteration of standard MSNs is described as ‘vesicular multilamellar mesoporous silica’ (VMMS). The reason for interest in VMMS structures, possessing three or more shells, is because these multi-shelled mesoporous silica nanospheres exhibit enhanced features, such as a prolonged release time of its contents and the ability for heterogenous catalysis. The circumvention of immediate burst release, often seen in stimuli-responsive nanocarriers, is paired with a decreased level of in vivo toxicity, a prolonged time interval in which the drug can exert its effect and prevention of immediate cell receptor saturation, which would otherwise deprive remaining drug molecules of any value. Zhang *et al.* previously reported the synthesis of VMMS nanospheres with adjustable number of layers by simply altering the ratio of structure-directing agents (9). The quaternary ammonium bromide co-surfactants didodecyldimethylammonium bromide (DDAB) and cetyltrimethylammonium bromide (CTAB) are utilized in cationic-cationic co-surfactant templating for the succeeding silica onion-like nanoparticles. Bundles of co-surfactant bilayers separated by an aqueous solvent, make up the lamellar phases and are ultimately replaced by mesoporous silica on both sides of the bilayers after calcination (Figure 1A). To ensure good dispersity, stabilization of coalescences caused by Brownian motion and the prevention of Ostwald ripening, the structure directing agent CTAB was used (10, 11). This co-surfactant is known for its long alkyl chain, which on itself imposes a protective effect, and is commonly used for silica and metallic nanoparticles preparation. An augmented packaging capacity and the ability for bilayer formation at lower co-surfactant concentrations was realized by addition of the double chain co-surfactant DDAB. Furthermore, the double-chained characteristic of this cationic amphiphile will ensure robustness in supramolecular assemblies, thus reinforcing the structural integrity of micelles (9). The use of co-surfactants over surfactants creates additional advantages because their relatively small polar head groups subsequently result in an increase of active surface area, a greater packaging parameter, decreased facial tension and a tendency to transform to worm-like micelles. Their long alkyl chains translate to a diminished critical micelle concentration (CMC) due to an increased hydrophobic character, and finally the repulsive interactions between both co-surfactants ensues lamellar phase formation at lower concentrations (12). Jiang *et al.* were able to modify the morphology of VMMS nanospheres by adding β-cyclodextrin (β-CD) as a regulator to the CTAB/DDAB dual-templating process (13). The spherical architecture of the vesicles was transformed into a worm-like micellar complex by merely increasing the concentration of β-CD. Furthermore, greater concentrations of β-CD resulted in the restructuring of worm-like micelles into spheroids. Additionally, the number of layers decreased with increasing amounts of β-CD during the templating process. The molecule cyclodextrin consists of six, seven or eight glucopyranose units ordered in a macrocyclic structure, called α-, β- and γ-cyclodextrin respectively (Figure 1B). The overall 3D morphology of cyclodextrin resembles that of a truncated cone with a polar exterior and a less polar interior. Similar to calixarenes or pillararenes, cyclodextrin is a host-molecule containing a hydrophobic cavity which is able to encapsulate hydrophobic guest-molecules in order to entrap them indefinitely or release them upon exposure of external stimuli. For example, Shouhui *et al.* took advantage of weakening interactions between the host-guest complex by raising hydrophilic properties of guest-molecules through different external stimuli exposure. In near-neutral pH conditions, benzimidazole (Bzl) has hydrophobic sites, thus making it able to bind with cyclodextrin. However, the binding constant between the host-guest complex decreases substantially when Bzl gets protonated under acidic conditions. The drug molecules azobenzene and ferrocene exhibit a similar solubility shift when exposed to different wavelengths and voltages, respectively (14). Between α-, β- and γ-cyclodextrin, the molecule β-CD is commonly used because of its remarkably low aqueous solubility compared to its α- and γ-counterparts. This ascribed to a high energy H-bond network between C2 and C3 hydroxyl groups, making it ideal for encapsulating hydrophobic guest molecules such as alkyl chains of surfactants. Interestingly, β-CD can interact with micellar structures by altering the micellar composition, thus modulating the properties and morphology of already formed vesicular structures (15). The favored entrapment of CTAB hydrophobic tails in β-CD cavities was speculated during testing of CTAB/DDAB structures, thereby explaining the shift of micellar morphology. CTAB/β-CD complexes exhibit a higher binding constant opposed to DDAB/β-CD complexes with experimental values of 5 × 10^4^ M^−1^ and 210 M^−1^, respectively (15, 16).

**Figure 1:**
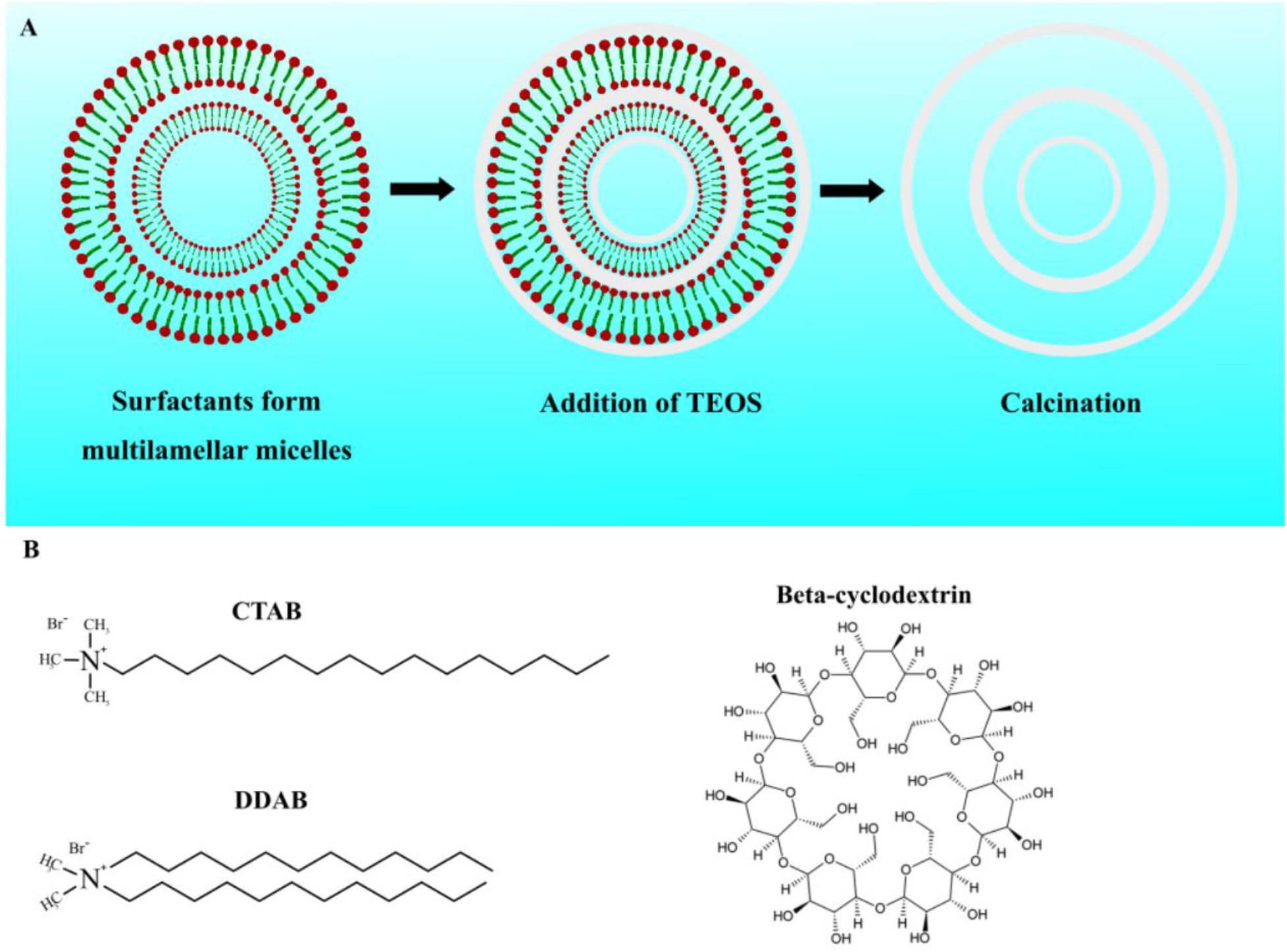
the templating process and materials for multilamellar mesoporous silica nanoparticles. (A) The templating process including multi-layered bilayer formation, TEOS disposition and calcination. (B) Chemical structures of CTAB, DDAB and beta-cyclodextrin.

The goal of this report is to analyze the differences and similarities between non-lamellar and multilamellar, spherical- and worm-like MSNs in order to determine their potential purposes in biomedical applications. Comparing volume, specific surface area, stability, drug-loading capacity, release kinetics and biocompatibility are solid starting points for characterization of both mesoporous silica micelles. Drug-packaging capacity and release kinetics were determined by loading the model drug aspirin into the VMMS- and worm-like multilamellar mesoporous silica (WMMS) structures. The small molecular dimensions of aspirin allow for easy infiltration and diffusion within the porous silica matrix.

## 2. EXPERIMENTAL PROCEDURES

### 2.1 Materials

Chemicals and solvents were purchased from commercial producers. Beta-cyclodextrin hydrate (99%) and didodecyldimethylammonium bromide (99%) were purchased from Acros Organics. Cetyltrimethylammonium bromide (≥98%) was purchased from Sigma Aldrich. Ammonia solution (32%) was purchased from Merck Milipore. Tetraethylorthosilicate (98%) was purchased from Alfa AesarTM. All chemicals were used without any further purification.

### 2.2 Synthesis of W- and VMMS nanoparticles

The dual-templating procedure including synthesis and modification, the disposition of silica and the removal of the cationic surfactants are elaborated in this paragraph. Both procedures were derived from Zhang *et al.* and Jiang *et al.*, respectively (9, 13). The co-surfactants CTAB and DDAB were dissolved in 70 ml MilliQ during continuous stirring at 30 °C. First, an amount of 0.284 g CTAB was added to the aqueous solution before introducing specified molar ratios of DDAB (CTAB:DDAB being 1:0.312, 1:0.625, and1:0.832). The mixture was stirred until both surfactants were completely dissolved. Further, 1.38 ml of ammonia was added as a Stöber-like synthesis catalysator and left stirring for 2h. Second, 4.29 ml of tetraethylorthosilicate (TEOS) was dissolved while stirring at 30 °C for 24 h. The mixture was subsequently transferred to a Teflon-lined autoclave and stored at 100°C for 24 h under static conditions. The product was collected using a centrifuge 3 times at 12000 rpm using a Eppendorf centrifuge 5804R. Finally, surfactants were removed by calcination using a tube furnace at 550 °C for 6 h. The different ratios were named VNMS, VMMS-1 and WMMS respectively. The transition of vesicular MSNs to worm-like MSNs using the regulator β-CD was planned to be analyzed after the synthesis of a β-CD/CTAB/DDAB system. First, 0.284 g CTAB and 0.224 g of DDAB were dissolved in 70 ml of MilliQ water and left stirring at 30 °C until the solution became clear. Two experimental amounts of β-CD (0.1 g and 0.2 g) were separately added to their own CTAB/DDAB solution and stirred until completely dissolved. The remaining products ammonia, TEOS were added in the same amounts as the previous experiments and stirred for 24 h. Again, the hydrothermal treatment with a Teflon autoclave at 100 °C for 24 h was performed. Products were centrifuged at … rpm (rotator …) and surfactants were removed by calcination using a tube furnace at 550 °C for a time period of 6 h. The samples were called β-VMMS and β-WMMS, respectively.

### 2.3 Characterization

Transmission electron microscopy imaging of W- and VMMS nanoparticles was done by a TECNAI G2 Spirit TWIN from FEI, operating at a 120 kV accelerating voltage. The W- and VMMS nanoparticles were dispersed in ethanol and dropcasted onto a copper grid and dried until all ethanol was evaporated. No additional staining was used for imaging. N2 adsorption/desorption experiments were performed using a Micrometric Tristar II. The specific surface areas were analyzed by the BET (Brunauer-Emmet-Teller) method and the pore size distribution was assessed by the BJH (Barret-Joyner-Halenda) method.

#### Important note

The following explanations regarding “Adsorption and release kinetics” and “Biodegradation” could not be realized due to unforeseen circumstances, which could not be resolved within the limited timespan of the research. Both sections were kept to illustrate the procedure, which normally would have taken place. A discussion based on literature studies has replaced the elaboration of the planned results.

### 2.4 Adsorption and release kinetics

The adsorption and release of the model drug aspirin in wormlike- and vesicular MSNs was assessed in the following paragraph. Both sets of nanoparticles (0.15 g) were exposed to aspirin in a 20 ml aspirin-ethanol solution (20 mg ml-1) at 25 °C. Subsequently, the mixture was stirred for 24h at room temperature. After incubation, the obtained solution was centrifuged at 6000 rpm for 10 minutes to retrieve the aspirin-loaded nanoparticles, which were dried at 25°C. The filtrate of the centrifuged mixture was diluted to 25 ml with ethanol and analyzed using an UV-VIS spectrophotometer at 275 nm. The amount of adsorbed drug was calculated by comparing mass before and after adsorption. Measurements of each sample were conducted three times such that an average result could be calculated. The drug loading content and encapsulation efficiency were estimated with the following calculations:

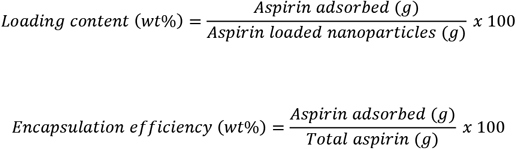

After successful loading of aspirin in the samples, the release profiles of their contents were assessed. An amount of 100 mg of the aspirin-loaded nanoparticles were dispersed in a 50 ml phosphate buffer solution (pH 7.4) and continuously stirred at 300 rpm and at a temperature of 37 °C. A sample of 3 ml was taken from the solution and replaced immediately by 3 ml of fresh phosphate buffer solution at each 30 min interval for 3 h and eventually after every 1 h until the benchmark of 10h was reached. The obtained samples were centrifuged at 6000 rpm for 10 minutes and analyzed using the Varian Cary 500 Scan spectrometer at 275 nm. Again, the measurements were repeated three times to obtain an averaged value. Aspirin release (wt%) was calculated as followed:

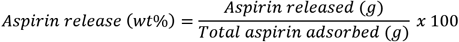

### 2.5 Biodegradation

Samples VNMS, VMMS-1, WMMS, β-VMMS and β-WMMS were each diluted in an 1 mg/ml aqueous solution using MiliQ and subsequently dialyzed against a NaCl solution (0.9%) using a dialysis tubing membrane, while a reference sample was made for each NaCl solution. Samples (1 ml) of the solution were taken at specific time-intervals and immediately refreshed by adding 1 ml of new NaCl solution. An amount of 0.3 ml of 1M hydrochloric and 1 ml ethanol was added to each sample before addition of β-silicomolybdate as a way to stabilize the complex. After an incubation time of 15 min, 1.5 ml of 5% ammonium molybdate tetrahydrate was mixed with the solution until the appearance became yellow. After 20 min, 1 ml of 1M oxalic acid was added and incubated for 10 min, such that 0.5 ml of 2% ascorbic acid could added to the mixture. Absorption was measured 10 min later at 810 nm using an UV-VIS spectrophotometer. The NaCl reference samples were subjected to the same processes.

## 3. RESULTS

### 3.1 Characterization

Non-lamellar mesoporous silica nanospheres were made to illustrate the effect of co-surfactant dual templating on the overall structure of the silica nanoparticles. The production of mesoporous silica nanoparticles (MSN) by a low CTAB/DDAB ratio system produced non-layered mesoporous silica structures, which are likewise able for aspirin uptake and release. However, the morphology diverges significantly from the MSNs made by higher ratios CTAB/DDAB of the cationic-cationic co-surfactant templating route. The non-lamellar and multilamellar MSNs were synthesized by adding the structure-directing agents CTAB and DDAB in proportions of 1:0.312, 1:0.625 and 1:1.832, thus making ordered lamellar formation of silica possible. A fourth sample, which was previously synthesized and analyzed, was compared to the other MSNs and had a ratio of 1:104 CTAB/DDAB, named VMMS-2. The difference between the low and higher ratios were apparent in TEM imaging (Figure 2). Moreover, certain distinctions between the three different synthesized multilamellar MSNs proved the utility of tuning the CTAB/DDAB ratio. The non-lamellar MSNs exhibited a mean diameter of 130 ± 25 nm, which implies an overall nano-scaled size but a considerable size distribution. The spherical structures are commonly referred to as spheroids, alluding to their oval nature instead of ideal orb-like features. This discrepancy in their spherical geometrical dimensions could explain the variance among the measured diameters, rather than the overall size distribution itself. A small decline in diameter was observed with increasing DDAB within the dual-templated MSNs regarding VMMS-1 (117 ± 12 nm) and WMMS (107 ± 36 nm). However, the multi-layered product VMMS-2 was characterized by remarkably large particles with a wide range of variable widths, indicating a clear diversion from the previous values. On basis of empirical data combined with visual subjective observation of VMMS-2, one must conclude the diameter variance to be heavily dependent on size distribution rather than geometrics. The biggest difference, only stated in theory this far, between the synthesis procedures is the emergence of lamella when utilizing higher ratios within the dual-templating approach and was clearly visualized by TEM imaging. The non-lamellar MSNs show an orderly structured complex of silica mesochannels. These worm-like pores are not sequentially divided in layers but form one hierarchical silica network uninterrupted and interconnected with one another. This mesh of random oriented mesochannels, although similarly capable for drug loading and release, is clearly distinct in morphology opposed to VMMS-1, WMMS and VMMS-2. A large number of lamellae are formed around a central core when the template is constructed with a 1:0.625 CTAB/DDAB ratio. The radial mesochannels are replaced by porous sheets, which are distinctly separated by aqueous solvent in a uniform manner. Apart from VMMS-1, the emergence of a clear core becomes distinguishable from the layered shell when DDAB plays more role in the structure of MSNs. The dark spotted centre has faded into the most translucent part of the MSN complex, thus illustrating the presence of silica-free spaces and the transformation of hierarchical structures to a more core-shelled morphology. The lamellae itself reduce drastically in numbers as the core grows due to greater amounts of DDAB in the shell. The double-chained co-surfactant seems to be responsible for the occurrence of both effects. The mere differences between CTAB and DDAB are alkyl length and the number of tails attached to the headgroup. Furthermore, a slight geometrical difference was noticed between the multilamellar MSNs. Both DLV-1 and DLV-3 samples contained exclusively mesoporous silica nanovesicles, characterized by the propensity of forming a more spherical architecture. However, DLV-2 showed among the nanovesicles a seemingly subpopulation of worm-like micelles, which are absent in the latter two MSNs and non-lamellar silica nanoparticles. Clearly, DDAB plays a big role in the morphology of these multilamellar MSNs by gradually overtaking the prominent role of CTAB as the structure-directing agent. Aparajita *et al.* observed the role of water loading in rod-like to spherical structure transitions of a DDAB/water/cyclohexane system. The dichotomy of hydration and dehydration within co-surfactant systems seems to be correlated to the structural changes induced by co-surfactant DDAB. The enlarged core offers more room for hydration, thus lessening the degree of dehydrated areas which would be within the shell. Interactions between surfactant head groups and water on their interface could impact the eventual shape of the micelles (17).

**Figure 2:**
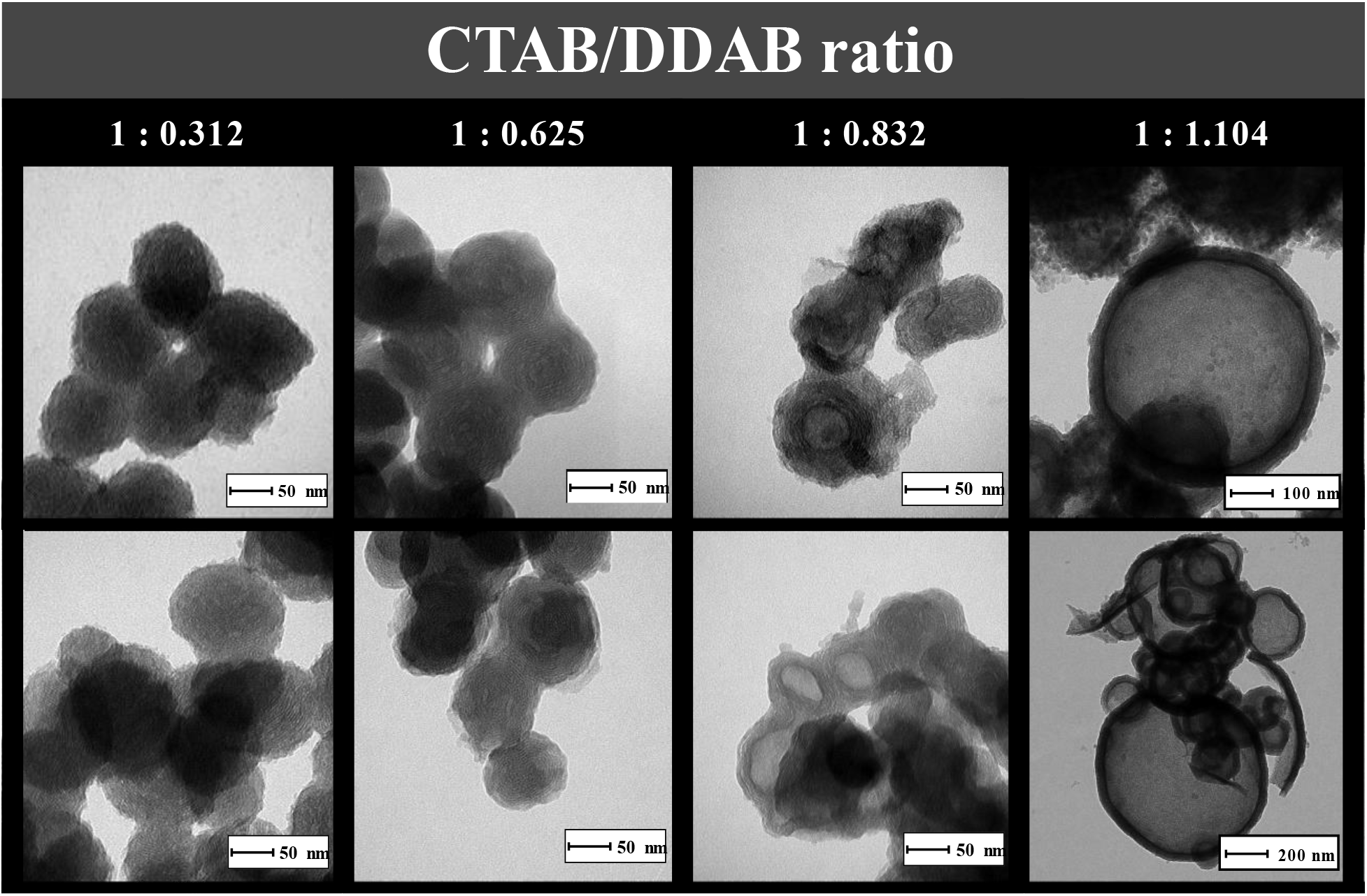
TEM imaging of MSNs using different CTAB/DDAB ratios. Two examples of MSNs of each ratio (1:0.312, 1:0:0.625, 1:0.832 and 1:1.104) are displayed using tomography emission microscopy (TEM) imaging.

The utility of multilamellar MSNs is determined by the degree of loading and release of therapeutically favorable drug molecules. Higher specific surface area (total surface area per unit of mass) correlates directly to the magnitude of potential physical adsorption of molecules on a solid surface. Ideal MSNs exhibit large specific surface areas to enclose as much drug molecules as possible. The model used for determining this feature presupposes the assumptions that inert gas like nitrogen (Henry’s law) at isothermal conditions solely physically adsorbs on solid silica layers and adjacent N2 layers, thus forming stacks of layers on top of a prior formed N2 monolayer. Furthermore, the uppermost layer exhibits equal adsorption and desorption rates, thus existing in an equilibrium. This extension of the Langmuir theory is known as the Brunauer-Emmet-Teller (BET) theory and is commonly used for determining specific surfaces areas of MSNs. DS, DLV-1 and 2 were analyzed by filling their pores with pressured nitrogen gas in order to determine the adsorbed amount of N2 and subsequently calculate the specific surface area of each MSN (Figure 3A). The non-lamellar MSNs proved to possess the smallest BET surface area with a value of 701.7 m^2^/g, followed by the multilamellar MSN DLV-2 with a value of 886,8 m²/g. TEM imaging previously confirmed DS particles to exceed DLV-2 in size, hence the formation of multiple lamellae must be responsible for the increase in bulk volume. The biggest specific surface area was spotted in DLV-1 (938,7 m^2^/g), which exhibits the most lamellae and a small aqueous core opposed to MSNs with larger CTAB/DDAB ratios. Also, the frequent formation of worm-like micelles could be the reason for the small difference between DLV-1 and -2, because of the superior volume in cylindrical structures compared to MSN spheres. Furthermore, all N2 adsorption-desorption isotherms of all examined samples share the same characteristics found in mesoporous structures, and is referred to as a type IV isotherm. The transition from the first trend to the first linear part in the graph, as seen at lower pressure differences, is an indication of monolayer coverage completion and the beginning of multilayer adsorption, also commonly found in type II graphs. The slim H4-type hysteresis loops are an allusion to the presence of narrow slit-like pores. Additionally, it is commonly present in isotherms of hollow spheres with walls composed of ordered mesoporous silica. Moreover, this hysteresis type is associated with capillary condensation at higher pressure differences. Nitrogen will condense to a liquid-like state in a pore if the pressure p is less than the vapor saturation pressure p° of the bulk liquid. Hysteresis due to capillary condensation only occurs when the pore diameter exceeds a critical width (4 nm diameter for N2 at 77K), which depends on the adsorption system and temperature (18). The cumulative volumes of pores between 1.7 nm and 300 nm width was determined for the same samples (DS, DLV-1 and -2) by applying the Barrett-Joyner-Halenda (BJH) model on the N2 adsorption-desorption isotherms (Figure 3B). Again, DLV-1 MSNs exhibited the best results with a cumulative value of 0,68 cm^³^/g, while DS nanoparticles had the lowest volume of 0,52 cm³/g and DLV-2 a volume of 0,63 cm³/g. No, substantial changes in cumulative volume between the multilamellar MSNs were observed, but did differ remarkably from non-lamellar MSNs as seen in sample DS. Desorption isotherms were used to analyse pore size distribution (PSD) because this state corresponds to a more stable condition of the pores. The PSD among the three samples seem fairly similar, while pores with a width of around 2.75 nm exhibit the highest pore volume. Most pores are above the 4 nm width critical threshold for capillary condensation to occur as mentioned before (24 datapoints are outside the graphs abscissa). However, different reports of BET and BJH analysis criticize the methods to be outdated and inaccurate when it comes to investigating either micropores or narrow mesopores. Both techniques are based on Kelvins equation (Young-Laplace), which consequently assumes that pore fluid has equal thermophysical properties as bulk fluids, yet both experimental and theoretical works have proven their thermodynamic properties to be considerably different. As a consequence, BET has been proven to be unreliable within a factor of at least 4 when it comes to surface area and pore volume analysis. Other argumentations, such as the complication of Henry’s laws assumptions, to devalue these traditional thermodynamic, macroscopic methods, are in abundance, thus one should be cautious when interpreting the obtained results acquired from both the BET and BJH method. Possible alternatives to validate the results is by the use of Density Functional Theory (DFT) or molecular simulation methods, like the Monte Carlo simulation method (MC). These methods provide a more realistic description of thermodynamic characteristics by connecting macroscopic properties with molecular behavior (19, 20).

**Figure 3:**
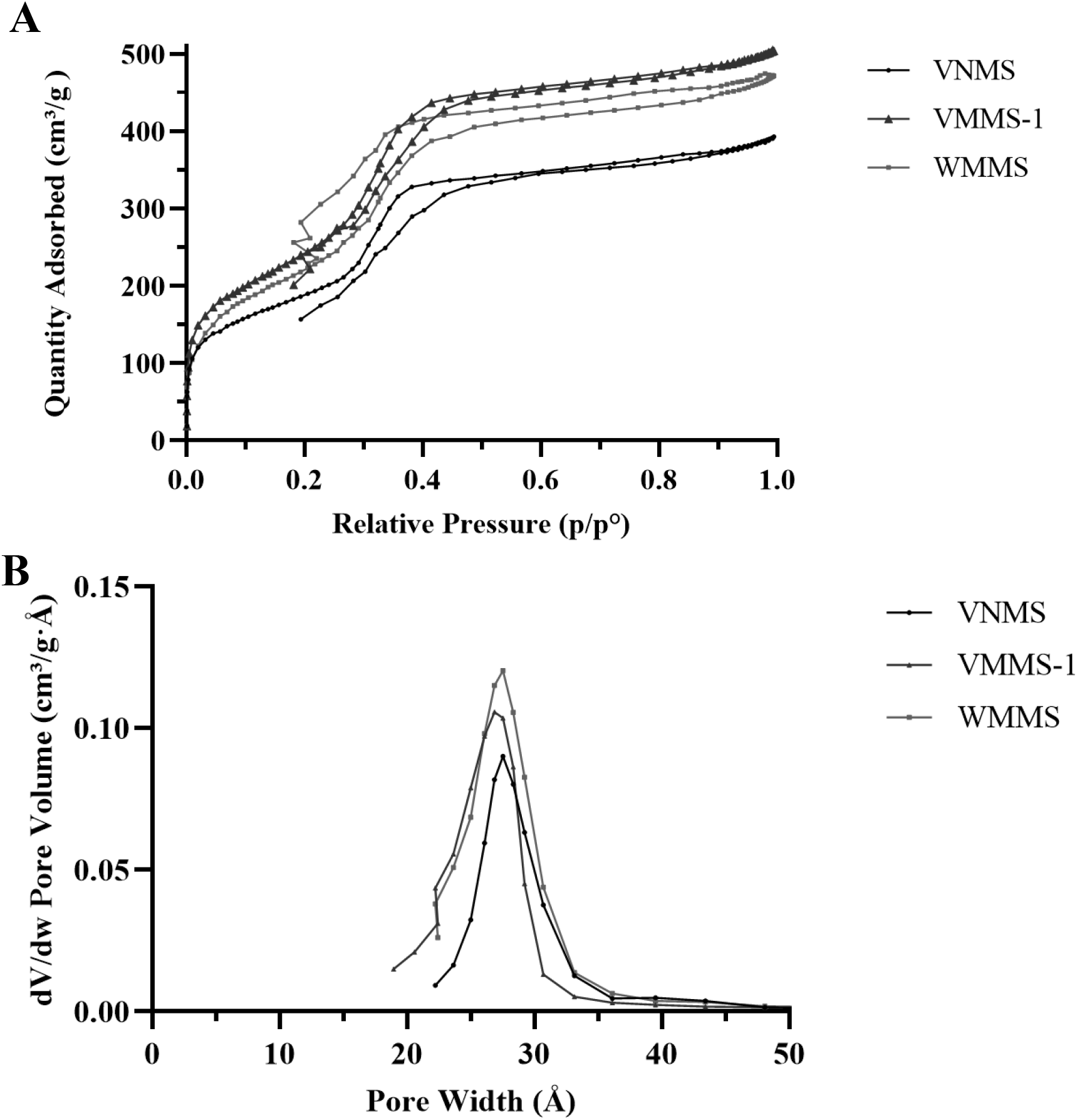
BET and BJH analysis of multilamellar mesoporous silica nanoparticles. N_2_-isotherms of VNMS (●), VMMS-1 (▲) and WMMS (■) using (A) the Brunauer-Emmet-Teller (BET) method and (B) the Barrett-Joyner-Halenda (BJH) method.

### 3.2 Adsorption & desorption kinetics

The shape, the extent of lamination and the functional surface coverage are all suspected to influence the degree of drug loading and release. The synthesis of the non-lamellar and multilamellar MSNs were derived from Zhang *et al.*, which conducted drug load- and release experiments for all MSNs with their respective CTAB/DDAB ratio (9). These results will be further discussed and compared to a follow-up study in this section. As mentioned before, β-CD can alter the overall morphology of vesicular and worm-like micelles by entrapment of said surfactants in a hydrophobic inner cavity. Additional research conducted by Jiang *et al.* utilized this particular feature of β-CD to investigate the drug loading- and release profiles of micelles correlated with their structure (13). Exposing vesicular micelles, made by the addition of 0.142 g CTAB and 0.112 g DDAB to 35 ml deionized water, to specific amounts of β-CD (0.05, 0.1, 0.15, 0.2 and 0.25 g) made excellent candidates to analyze the previous hypothesized correlation. This report, will focus mainly on the solutions with 0.05 g and 0.1 g β-CD, for these were similarly replicated as seen in section 2.0 ‘Materials & Methods’. Moreover, these amounts do not affect solely the number of layers, but also induce the transition from vesicular to worm-like micelles.

First, the drug loading capacity of DS, DLV-1, DLV-2 and DLV-3 were determined and exhibited values of 35.5 wt%, 28.7 wt%, 54.1 wt% and 40.8 wt%, respectively. The multi-layered MSNs with added β-CD (0.05 and 0.1 g) accommodated 29.1% and 45.7% of drugs. Both release studies cannot be compared, considering that these studies used different drug models. However, these reports did agree that the amount of drug loading was dependent on the cavity size of vesicles. This said, DLV-1 had a greater specific surface area opposed to DLV-2, so other factors than surface area must come in to play. The altered MSN vesicles with β-CD showed a significant increase in drug loading by 16.6 wt%. The transition to a worm-like state combined with a decreased number of layers had a major influence on the potential of MSN to accommodate therapeutic drugs. DLV-2 was earlier characterized with a subpopulation of worm-like vesicles and an overall decrease in lamellae, thus possibly explaining their superior capacity. The two aforementioned studies were completer with additional release profile analysis with again each their own model drug. The general conclusion was a relatively steady release rate and a prolonged release time associated with the number of layers. Sample DS exhibited a continuous rapid release of its contents opposed to the multi-layered MSNs. The more layers an MSN possessed, the more time was needed to reach a total drug content depletion of the nanoparticle. The initial rapid release of all samples could be due to drug clogging in the outer pores of the structure. Thereafter, all drug contents need to transverse a spatial-dependent amount of layers obstructing them for release in the outlying matrix. This was became apparent when MSNs with less lamellae required less time to release all their contents. Jiang *et al.* obtained similar conclusions between the vesicular- and worm-like micelles, with the latter having less layers than the former. Apart from tuning the number of layers in order to retard the release time of drug loaded MSNs, other methods have been reported to control the release rate. Chemical modification of the mesoporous silica surface allows a broad variety of hydroxyl functionality, thus tailoring the surface area for specific needs. Aspirin was not only chosen because of its small molecular dimensions, but also because of the complementary physical mechanisms between the drug and bare silica. The carboxylic acid group of aspirin interacts readily with the hydroxyl groups of silica. The hydrogen bonding promotes aspirin physisorption, meaning the force of attraction is reversible and allows the formation of multi-molecular layers. These features could be enhanced by post-grafting modification of mesoporous silica with carboxyl- and amino groups. Research group Gao *et al*. used feeble chemical interactions between silanol and 3-aminopropyltriethoxysilane (N-TES) or 3-(2-aminoethylamino) propyltrimethoxysilane (NN-TES), while Xu *et al*. trimethylsilyl-carboxyl groups to regulate the release of drug molecule famotidine (21, 22). Consequently, the adsorption capacity of the surface-modified MSNs was vastly improved and deceleration of drug release from the pores of mesoporous silica was achieved. An interesting addition to drug-loading and release analysis would be the comparison between their kinetic profiles and the cumulative volume of their pores with a specific width range (1.7 – 300 nm in this study). Diffusion of drug molecules is largely dependent on the intrinsic mobility of the drug molecules within the pore, which becomes greater with an increasing volume.

### 3.3 Biodegradability

These multilamellar MSNs are ultimately made for various biomedical objectives, which requires administration of the nanoparticles in human body. However, accumulation of these MSNs within tissue could cause adverse effects, such as immune responses. Therefore, nanoparticles are made of material susceptible to external factors inducing biodegradability of the structure. Silica is an excellent material for nanoparticle synthesis because of biocompatibility, post-grafting functionalization, inertness under many conditions, easily to form structured particles and biodegradability. The dissolution of silica consists of three parts: (1) water is adsorbed into the siloxane mesh, (2) siloxane is gradually hydrolysed into silanols and (3) an ion-exchange process, which uses nucleophilic attacks of hydroxyl groups, leads eventually to the leaching of silicic acid (23). The end products diffuse through the bloodstream of the lymph lobe to be eventually cleared in urine. The hydration-hydrolysis-ion-exchange process can be replaced by NaCl biodegradation of siloxane to easily measure the degree of biodegradability of MSNs. Administration of medicines commonly contains saline solutions, which are mimicked with these NaCl dispersions. The chloride ions within the solution is able to perform a nucleophilic attack on siloxane, consequently repositioning the groups and therefore making them more susceptible to hydrolysis. Both methods subsequently result in dissolution of the outer surfaces, working their way to more internal structures (24). Multilamellar MSNs will require a delicate balance between the degree of biodegradability and the timespan required for drug release to completion. This type of nanoparticles need to withstand the physiochemical, environmental influences long enough to effectively discharge therapeutic drug molecules over a well-defined timespan. Shortly after complete desorption of the drug, the structural integrity of the MSNs should break down to their initial building blocks in order to be eliminated from the body. This will require fine-tuning of the particles chemical structure and size to obtain perfect timing between both elements. The use of silica hybrids will fasten the dissolution of silica within certain types of environments. Incorporation of redox-sensitive bridging groups within silica, such as disulphide bridges will make them more susceptible for glutathione-rich environments as seen in tumours of inflamed tissue. Enhanced hydrolytic degradability within the framework is achieved by different approaches, making these structures pH-sensitive. The inclusion of redox-, pH-, enzymatic- or biochelation-mediated lysis mechanisms does not only accelerate the dissolution process, but additionally adds a targeting-mechanism by solely allowing fastened degradation if specific external stimuli are present (25). Apart from alterations of the chemical built, the number of layers of the multi-layered MSN structure will likewise impact the time until full biodegradation has occurred. This feature should also be taken into account when tuning the degree of dissolution of multilamellar MSNs.

## 4. CONCLUSION

In conclusion, this report describes a customizable system where the number of lamellae of multilamellar mesoporous silica nanovesicles could easily be adjusted and additionally resulted in a prolonged release of adsorbed drug molecules. A heightened CTAB/DDAB ratio appeared to decrease the amount of layers once the multilamellar morphology had formed, increased the pore volume, increased the drug-loading once a clear core had formed and both the specific surface area and prolonged drug release seemed to be layer-dependent. The ratio of 1:0.832 appeared to be the most balanced ratio of all four, which resulted in a multilamellar core-shell morphology, a high specific surface area, relative large pore volume, superior drug loading and a prolonged drug release, thus being an excellent candidate for further improvements. Some potential future outlooks could examine the effect of beta-cyclodextrin on the multilamellar MSNs, such that one is able to tune the templates morphology if the structure is already formed. Also, to prevent the accumulation of MSNs using biodegradability, the tests proposed in the section ‘Biodegradability’ could be conducted, while also trying stimuli-responsive groups to promote the dissolution. Post-grafting possibilities could explored to enhance the drug adsorption of the multi-shelled silica nanosphere. These future studies, in combination with the previous obtained results, could provide a clear view on the potential applications for these multilamellar mesoporous silica nanoparticles.

## Acknowledgements

Both TF and NG are grateful for UV and AH to have granted us this opportunity to participate in their research project. TF thanks NG, UV for their contribution during the conducted experiments..

## Author contributions

UV and AH conceived and designed the research. UV, NG and TF performed synthesis and data analysis. UV performed all experiments. TF and UV carefully edited the manuscript.

